# Functional dissection of *SPOP* on the amino acid level reveals a comprehensive functional landscape of variants during tumorigenesis

**DOI:** 10.1101/2025.08.20.671394

**Authors:** Seong Kyoon Park, Jeongha Lee, Seon Ju Park, Ye Na Kim, Gi Hyun Shin, Kisoon Dan, Hee-Jung Choi, Dohyun Han, Byung Joon Hwang, Murim Choi

## Abstract

Numerous proteins display pleiotropic functions in different clinical contexts. However, molecular mechanism underlying such effects is rarely understood. Speckle-type POZ protein (*SPOP*) is a typical example, exhibiting tumor-suppressing or -promoting effects in different tumor types in accordance with different amino acid changes; specifically, two distinct sets of variants in *SPOP* are commonly found in subsets of prostate cancer and endometrial cancer patients. To comprehensively characterize the functional landscape of *SPOP* alteration, we performed a deep mutational screening (DMS), elucidating the functionality of 7,933 out of 8,228 possible single amino acid changes (96.4% coverage). Leveraging the observation that overexpression of human *SPOP* leads to yeast growth arrest, we assessed the functionality of each variant using a yeast proliferation assay. In addition, our approach combined long-read and short-read sequencing. Finally, our DMS model enables a clear distinction of likely-loss-of-function (LoF) variants that are enriched in prostate cancers and reveals their differential characteristics in both protein structure and genetic assessments. These results demonstrate the utility of our approach in high resolution mapping and amino-acid-level interpretation of protein function.

**Significance statement:** Genetic mutations often play different roles in cancer, driving or suppressing tumor growth depending on their molecular context. The protein SPOP is a striking example that acts as either a tumor-suppressor or oncogene depending on the specific mutation and tissue. However, understanding how thousands of possible mutations alter its function has been a major challenge. Here, we applied deep mutational scanning to experimentally measure the functional effects of nearly every possible amino acid change in *SPOP*. This work provides the first comprehensive functional map of *SPOP* variants, not only advancing basic knowledge of cancer biology but also establishing a framework for interpreting patient mutations in precision medicine.

## Introduction

As large-scale efforts to sequence broader populations and diverse disease cohorts have become increasingly common, the identification of variants of uncertain significance (VUS) has likewise ballooned, leading to an urgent need for accurate functional interpretation of these variants. Conventional approaches–leveraging evolutionary conservation (1,2) and machine learning models (3–6)–are frequently employed to predict the potential impact of VUS. However, these *in silico* predictions often lack biological validation and can suffer from inconsistent results. While human population frequency data serves as a robust line of evidence (7,8), its utility is often limited for rare variants and can be less informative for individuals of non-European ancestries due to data underrepresentation. In that context, deep mutational scanning (DMS) offers a high-throughput experimental framework to assign functional scores to artificially introduced variants within a specific cellular context (9–13). Indeed, a number of genes with disease association have been analyzed through this method to provide insights into the function of pathogenic variants (14–16) and sensitivity to drugs (17,18). Notably, the reliability of DMS depends heavily on the construction of a comprehensive and uniform variant library, as well as the establishment of assay systems that properly report on the gene’s function.

Interpreting the functional impact of genetic variants is particularly critical in the context of tumorigenesis, as progressive accumulation of genetic alterations is one of the fundamental mechanisms through which cancer cells gain a survival advantage. Distinguishing driver mutations from passenger mutations among these acquired variants is essential for elucidating tumorigenic mechanisms and informing clinical strategies. Notably, some cancer-associated genes such as *NOTCH1*, *TP53*, and *MYC* exhibit both tumor-suppressive and oncogenic features, whereby both loss-of-function (LoF) and gain-of-function (GoF) mutations can promote tumorigenesis in a context-dependent manner (19–21). It is especially crucial to dissect the molecular mechanisms by which distinct mutations in these bivalent genes contribute to cancer. Indeed, recent deep mutational scans of *CARD11*, *PTPN11*, and *DDX3X* effectively stratified variants into LoF and GoF classes, linking them to differential pathologies or caner types (10,22,23).

The Speckle-type POZ protein (SPOP) functions as a substrate adapter for the CUL3-RING ubiquitin ligase (CRL3) complex, facilitating the ubiquitination and proteasomal degradation of a broad range of substates, including proto-oncogenes and the bromodomain and extra-terminal domain (BET) proteins. As such, the interaction between SPOP, CUL3, and its substrates is essential for effective proteolysis (24). Beyond its role in the CRL3 complex, the functionality of SPOP is also determined by its structural properties, as SPOP forms a higher-order oligomer structure through its BTB and BACK domains. This oligomerization is crucial for modulating its ubiquitination efficiency through subcellular localization and stability modulation (25,26). Genetic variants altering any of these properties are known to contribute to tumorigenesis by activating malignant transcriptional mechanisms (27,28), including genomic instability (29,30), or modulating *SPOP* function through aberrant assembly (31). According to The Cancer Genome Atlas (TCGA), *SPOP* variants have been identified in a number of cancer types; the most notable are prostate adenocarcinoma (PRAD: 53 carriers/496 patients = 10.7%) and uterine corpus endometrial carcinoma (UCEC: 53/512 = 10.4%), with instances also recorded in cancers of the penis, liver, intestine, and urinary tract (32). While *SPOP* is widely regarded as a tumor suppressor in that its LoF variants lead to increased oncogenesis, certain mutations found in endometrial cancer have been reported to act as GoF, enhancing ubiquitination activity (33). These divergent effects have been examined in the context of differential mutation hotspots observed between prostate and endometrial cancers (25,34), and numerous studies have been conducted on the mechanism of *SPOP*-related tumorigenesis and the distinct functional impacts of its mutations; nonetheless, a curated variant profile for *SPOP* is still lacking, as evidenced by the absence of any entry in ClinGen and the presence of only five somatic mutations reported in ClinVar.

To generate a comprehensive functional map of *SPOP*, we applied a DMS approach to systematically assign fitness scores to individual variants. Leveraging our observation that overexpression of *SPOP* arrests yeast cell proliferation, we utilized a yeast-based growth assay system to dissect the functionality of individual amino acid changes. We also implemented a long-read sequencing-based analytic system and compared its performance to that of a short-read system. Our approach enables the functional characterization of genetic variants and facilitates the integration of mutation profiles from prostate and endometrial cancers with their corresponding functional impacts.

## Materials and Methods

### Yeast strain

Yeast assays used *Saccharomyces cerevisiae* strain JC-993 (MATa gal4 gal80 trp1-901 ura3-52 leu2-3 leu2-112 his3 URA3::GAL1-LacZ) with three constructs integrated via CRISPR-Cas9: pADH1(cr)-LexA-ER-B112 (35), activated by ß-estradiol to drive I-Sce I expression through the LexA promoter (top construct in SI Appendix, Fig. S1A); pADH1(cr)-Zif268-PR-MSN2, activated by progesterone to drive DMS library expression through the pZ promoter (36); and the landing pad construct, integration site for the DMS library (the second construct in SI Appendix, Fig. S1A). This construct was inserted between TRP5 and CWH41 loci on chromosome VII, and contains a dummy sequence under pZ, I-SceI under LexA, and KanR, conferring G418 resistance.

### Library design, cloning, and transformation

For DMS library construction, *SPOP* cDNA (CCDS11551.1) was divided into 16 overlapping tiles (54-122 bp) based on the availability of internal restriction enzyme (RE) sites. As high-quality oligonucleotide synthesis is length-limited (∼190 nucleotides), we introduced synonymous mutations when no suitable RE site was available. Within each tile, saturation mutagenesis was performed at the amino acid level by introducing 19 different substitutions, more than one synonymous changes, one in-frame deletion, and one stop-gain mutation per codon (37). All designed sequences were synthesized as oligonucleotides (Dataset S1) by Twist Bioscience (South San Francisco, CA). Oligonucleotides from different tiles were separated and amplified using tile specific PCR primers (Dataset S2). PCR was performed with 16 ng of oligonucleotide pool as template using Phusion High Fidelity DNA Polymerase (M0503, New England Biolabs, Ipswich, MA) for 18 cycles. To reconstruct the full-length *SPOP* cDNA, 5’ and 3’ flanking open reading frame (ORF) sequences corresponding to each tile were synthesized and sequentially ligated via compatible RE sites (SI Appendix, Fig. S1B). Completed library constructs were cloned into a vector backbone compatible with the landing pad (SI Appendix, Fig. S1C). To facilitate homologous recombination, the cDNA was flanked by homology arms corresponding to the 5’ (pZ promoter) and 3’ (KanR) regions of the landing pad cleavage site (the two construct shown at the bottom of SI Appendix, Fig. S1A).

Yeast cells were transformed through the LiAC/ss carrier DNA/polyethylene glycol method (38). The inducer ß-estradiol (E2758, Sigma-Aldrich, Saint Louis, MO) was applied to activate expression of I-SceI, which recognized and cleaved its RE sites flanking the pZ promoter and KanR, generating double-strand breaks for integration of the transformed library via homologous recombination (SI Appendix, Fig. S1B) (39–41). Yeast clones harboring the library were selected by supplemention of culturing media with 200 ug/ml of G418 (11-811-023, Thermo Fisher Scientific).

### Library screening

To identify differential enrichment of variants in the yeast, two separate pools were prepared containing 2.0 x 10^8^ cells each. Both were initially cultured in 50 ml of YPAD media supplemented with 200 ug/ml G418, shaked at 230 rpm for one hour at 30 °C. Afterwards, the media were replaced with 200 ml YPAD containing 200 ug/ml G418. For the experimental condition (the ‘treated’, *SPOP*-expressed pool), 500 nM progesterone (P8783, Sigma-Aldrich, Saint Louis, MO) was additionally supplemented; the other, untreated pool served as a control (non-expressed pool). Both cultures were then incubated under 3D culture conditions for 24 hours at 30 °C (42).

### Library sequencing

Genomic DNA was extracted from 1.0 x 10^8^ cells via the YeaStar Genomic DNA kit (D2002, ZYMO research, Irvine, CA). Integrated *SPOP* mutant constructs were enriched through 18 cycles of PCR amplification using 500 ng of gDNA as a template, primers flanking the *SPOP* construct (forward: 5’-AGCTGCATAACCACTTTAAC-3’, reverse: 5’- GCAAATTAAAGCCTTCGAGC-3’), and Phusion HF polymerase (M0503, New England Biolabs). Amplified products were pooled and purified through ethanol precipitation, agarose gel extraction, and ethanol precipitation. For short-read sequencing, library DNA was prepared using the TruSeq Nano DNA Sample Prep Kit and sequenced 2x300 bp on an Element FreeStyle AVITI. For long-read sequencing, 200 pmole of PCR product was sequenced using Ligation sequencing amplicons V14 (SQK-LSK114, Oxford Nanopore Technologies, Oxford, UK) via the MinION Mk1B platform with a R10.4.1 flowcell (FLO-MIN114, Oxford Nanopore Technologies). Base calling was performed in super-accurate mode with the *SPOP* wild type sequence as reference.

### Calculation of z-scores

Scores from short-read and long-read sequencing platforms were calculated separately. Short reads were preprocessed (adapter trimming and contaminant removal) using BBDuk with adapter and PhiX sequences provided, error-corrected with BBMerge, and aligned to the *SPOP* cDNA reference using BBMap. BBDuk, and BBMerge are from BBTools (43). Variants were called using AnalyzeSaturationMutagenesis in GATK (44), and the resulting *variantCounts* files used as per-variant count matrices. These matrices served as input for DESeq2 (45), and the enrichment ratio between *SPOP*-expressed and non-expressed pools was calculated by adapting a pipeline from a previous study (46). Before generating the DESeqDataSet, variants were restricted to only single variants and those with raw count ≥5. Two biological replicates were used and adjusted during DESeqDataSet generation. Since only LoF variants in the non-expressed pool were expected to remain detectable in the *SPOP*-expressed pool, size factors were estimated from stop-gain variants. Prior to DESeq2 differential enrichment analysis and z-score calculation, we removed variants with inconsistent detection across biological replicates.

Specifically, variants were retained only if they had non-zero counts in both replicates in the untreated samples. This pre-filtering step reduces stochastic artifacts driven by replicate dropouts. Then, a revised input matrix was generated and summary statistics obtained from DESeq. Finally, z-scores were calculated as the log 2 fold-change divided by its standard error.

Long-read data was limited to high-quality (‘PASS’) reads. To ensure accurate variant calling, hybrid error correction employing AVITI short-reads was performed using the *ratatosk* correct function with options -v -c 16 -G -Q 90 (47). Error-corrected long-reads were subsequently filtered by length using *Seqkit* (48), and regions outside the ORF were trimmed using *cutadapt* (49). Filtered reads were aligned to *SPOP* cDNA using *minimap2* with the -ax map-ont option (50), and variants were called from the aligned bam files by GATK AnalyzeSaturationMutagenesis. The resulting variant count matrix was processed as for short-reads, including the same filtering criteria, application of size factors, and generation of z-scores. Lastly, scores from short-reads and long-reads were combined using inverse-variance weighted average values. For variants only in short-reads, DESeq2 summary statistics and z-scores were used as-is. Two-tailed *P*-values were calculated with merged scores and adjusted by the Benjamini-Hochberg FDR method.

### Serial dilution-spotting assay

Expression plasmids carried the *GAL1* promoter for galactose-inducible expression and a LEU2 auxotrophic marker for plasmid selection. In clone testing, both CEN and 2μ origins were used for clones shown in in SI Appendix, Fig. S2, but only CEN for the experiment in Fig. 1B. Yeast strains individually transformed with the respective mutations were cultivated in SD/-Leu medium and diluted with fresh SD/–Leu medium to an OD of 1.0 (∼1 × 10 cells/mL).

**Figure 1.**
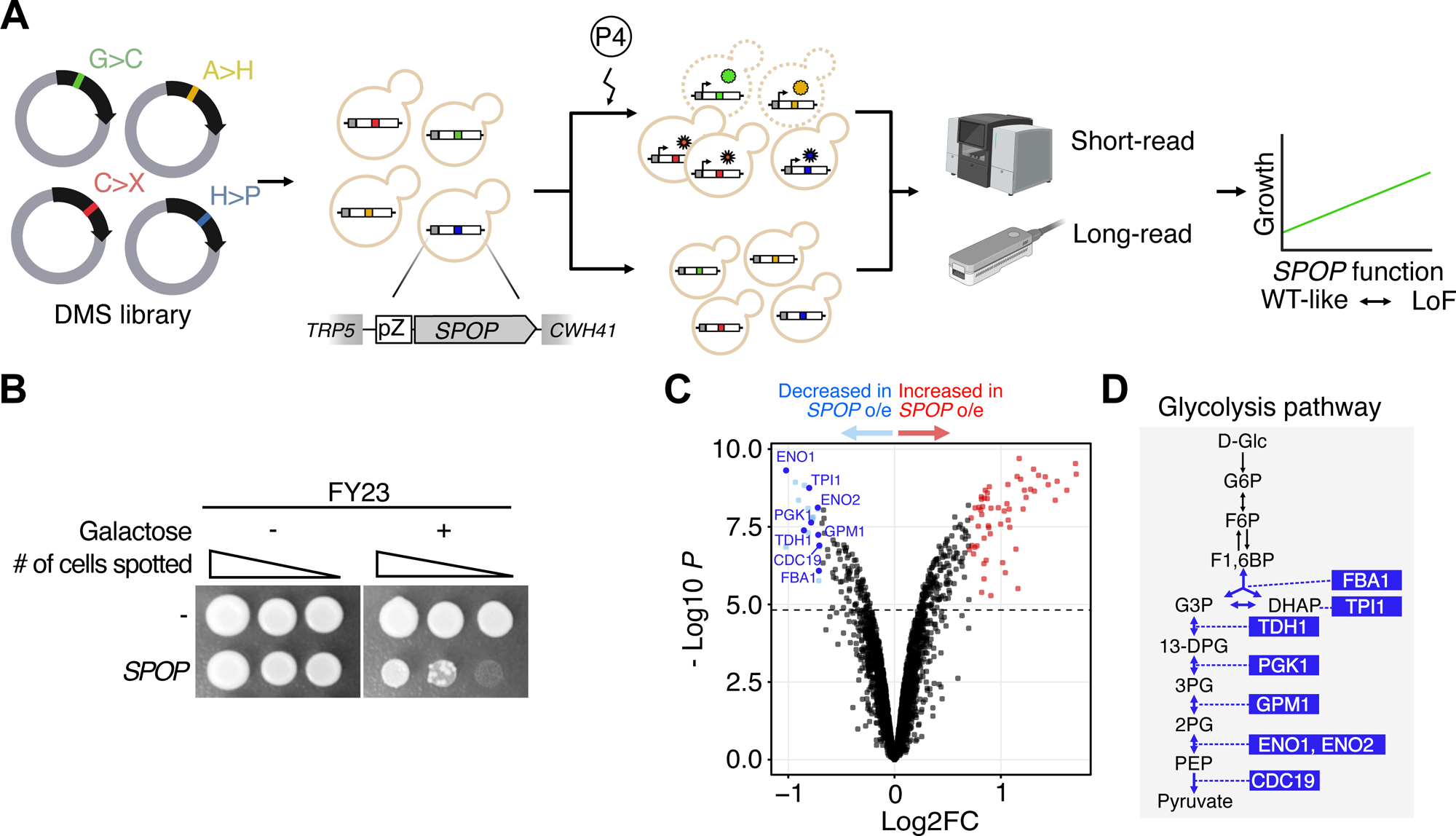
Overall scheme. (A) A DMS library was transfected into yeast. Expression of the introduced *SPOP* variants was induced by progesterone. After induction, genomic DNA was extracted from the surviving yeast cells, and the variants were identified using short- or long-read sequencing. The fitness score of each variant was based on the enrichment ratio in the *SPOP*-expressed pool compared to the non-expressed pool. (B) Effect of galactose-inducible *SPOP* expression on growth of the yeast strain FY23. Cells transformed with CEN plasmids were spotted onto plates containing galactose to induce overexpression. (C) Volcano plot of proteins differentially expressed upon overexpression of wild-type *SPOP* . (D) Position of highlighted proteins in the simplified glycolysis pathway. Significantly down-regulated enzymes were highlighted with blue. Abbreviations: P4, progesterone; D-Glc, D-glucopyranose; G6P, D-glucopyranose 6-phosphate; F6P, ß-D-fructofuranose 6-phosphate; F1,6BP, ß-D-fructose 1,6- bisphosphate; G3P, D-glyceraldehyde 3-phophate; DHAP, dihydroxyacetone phosphate; 13- DPG, 3-phospho-D-glyceroyl-phosphate; 3PG, 3-phospho-D-glycerate; 2PG, 2-phospho-D- glycerate; PEP, phosphoenolpyruvate.

Serial ten-fold dilutions (10^7^-10^3^ cells/mL) were prepared for each strain, and 5 μL of each dilution was spotted onto selective induction plates supplemented with 2% galactose.

### Yeast proteome analysis

Yeast expressing wild-type *SPOP* were cultured in YPAD media with and without 500 nM progesterone at 30 °C for one hour. Cells (1.0 x 10^6^) were harvested, lysed in SDS buffer via sonication (Branson Ultrasonics, Brookfield, CT), boiled, and lysate protein concentrations measured by tryptophan fluorescence assay (51,52). Preparation and analysis of peptides were described previously (53). Briefly, a total of 100 μg protein per sample was retrieved and resulting peptides were eluted, quantified, and labeled with TMTsixplex (Thermo Fisher Scientific) tag 126-128 for non-expressed or 129-131 for *SPOP*-expressed. The total of 96 fractions were concatenated into 24 and subjected to liquid chromatography-mass spectrometry (LC-MS). The resulting raw MS files were processed using Proteome Discoverer 3.1 (Thermo Scientific) against the *S. cerevisiae* protein sequence database (UniProt, July 2025). The FDR was set to <1%, and a co-isolation threshold of 50% was used for peptide quantification.

Differential expressed protein (DEP) analysis utilized the DEP package (54), with criteria of absolute log2 fold-change >0.7 and adjusted *P*-value <0.05. Gene Ontology enrichment analysis utilized the clusterProfiler package (55).

### Assigning variant tiers from platform-specific and merged scores

Variants observed in each sequencing platform were independently stratified by score; those with a Log2FC > -1 and adjusted *P*-value > 0.1, indicating no depletion in the *SPOP*-expressed pool, were classified as ‘Likely LoF’. Platform-specific tiers were used only for comparing the models derived from each sequencing platform.

Subsequent analyses employed merged scores, which were also stratified independently. Variants with Log2FC > -1 and adjusted *P*-value > 0.1, which did not significantly impact cell number or produced accelerated growth, were collectively classified as ‘Likely LoF’ due to failing to induce growth inhibition. And other remaining variants displaying depletion were classified as ‘Tolerated’.

### Quantification of mutant-harboring transcripts

Mutant-harboring transcripts were quantified by bulk RNA sequencing of *SPOP*-expression and non-expressing yeast pools, in three technical replicates. Total RNA was prepared using the TruSeq Stranded Total RNA Sample Prep Kit with Ribo-zero H/M/R (Illumina, San Diego, CA) and sequenced 2x150 bp on a NovaSeq X Plus (Illumina). Raw FASTQ files were aligned by STAR aligner in two-pass mode (56) to the *S. cerevisiae* reference genome (assembly R64), with wild-type *SPOP* appended. Mutation-associated expression was estimated by using pileup to stratify aligned reads by codon along the ORF and counting altered codons. Count matrices for each library were normalized by total read counts, normalized counts averaged within each pool, and differential transcript abundance assessed in terms of the log10-transformed ratio of pool means. Since a majority of assessed variants did not deviate from the mean, variants with normalized count ratio >1 were considered upregulated by the mutation, and those with ratio <-1 downregulated.

### Reference set-based evaluation of pathogenicity/benignity prediction

To calculate AUC values and likelihood ratios, we defined reference sets of benign and pathogenic variants. ClinVar variants annotated as *Pathogenic* or *Likely Pathogenic* were used as the pathogenic reference set and were stratified into somatic and germline subsets based on ClinVar classification type. Each ‘ClinVar-Somatic’ and ‘ClinVar-Germline’ set were composed with 4 and 10 variants. To increase power for somatic benchmarking, we additionally compiled a COSMIC somatic pathogenic reference set consisting of recurrent *SPOP* variants (occurrence count > 1), focusing on prostate cancer (*n* = 32). A curated germline LoF subset associated with Nabais Sa-de Vries syndrome was assembled from Nabais Sa *et al.* (57) (*n* = 4). As a benign reference set, we used missense variants observed in gnomAD (v4.1.0) (*n* = 60).

For each reference set comparison, we evaluated the binary classification to distinguish the pathogenic set from the benign reference set. Positive/negative likelihood ratio (LR+/-) was computed from sensitivity and specificity at the ‘Likely LoF cutoff’ (LR+ = sensitivity / (1 − specificity), LR- = (1-sensitivity)/specificity). AUC was calculated from the corresponding ROC curve using the continuous score, leveraging pROC R package.

### Comparison with external datasets

We utilized diverse *in silico* scores and cohort data in assessing our scoring system. Relative conservation scores were obtained from Consurf-DB (58), which provides pre-calculated evolutionary conservation profiles for Chain H of the 8DWV PDB structure. Conservation grades and solvent accessibility annotations (‘B/E’ for buried/exposed and ‘F/S’ for functional/structural) were used to evaluate tier-level enrichment of conserved or buried/exposed residues. *In silico* variant effect scores were obtained from AlphaMissense (4), EVE (5), and ESM1b (6). Variants annotation was performed using public databases, including gnomAD (v4.1.0; accessed on 12/02/2025), ClinVar (accessed on 12/01/2025), and COSMIC (accessed on 12/02/2025) (59). Structural and positional visualization utilized PyMOL with the 8DWV (for Fig. 2E, Fig. 4C-E, and SI Appendix Fig. S6, Fig. S8) and 3HQI (for Fig. 3F) PDB structures as templates (60).

**Figure 2.**
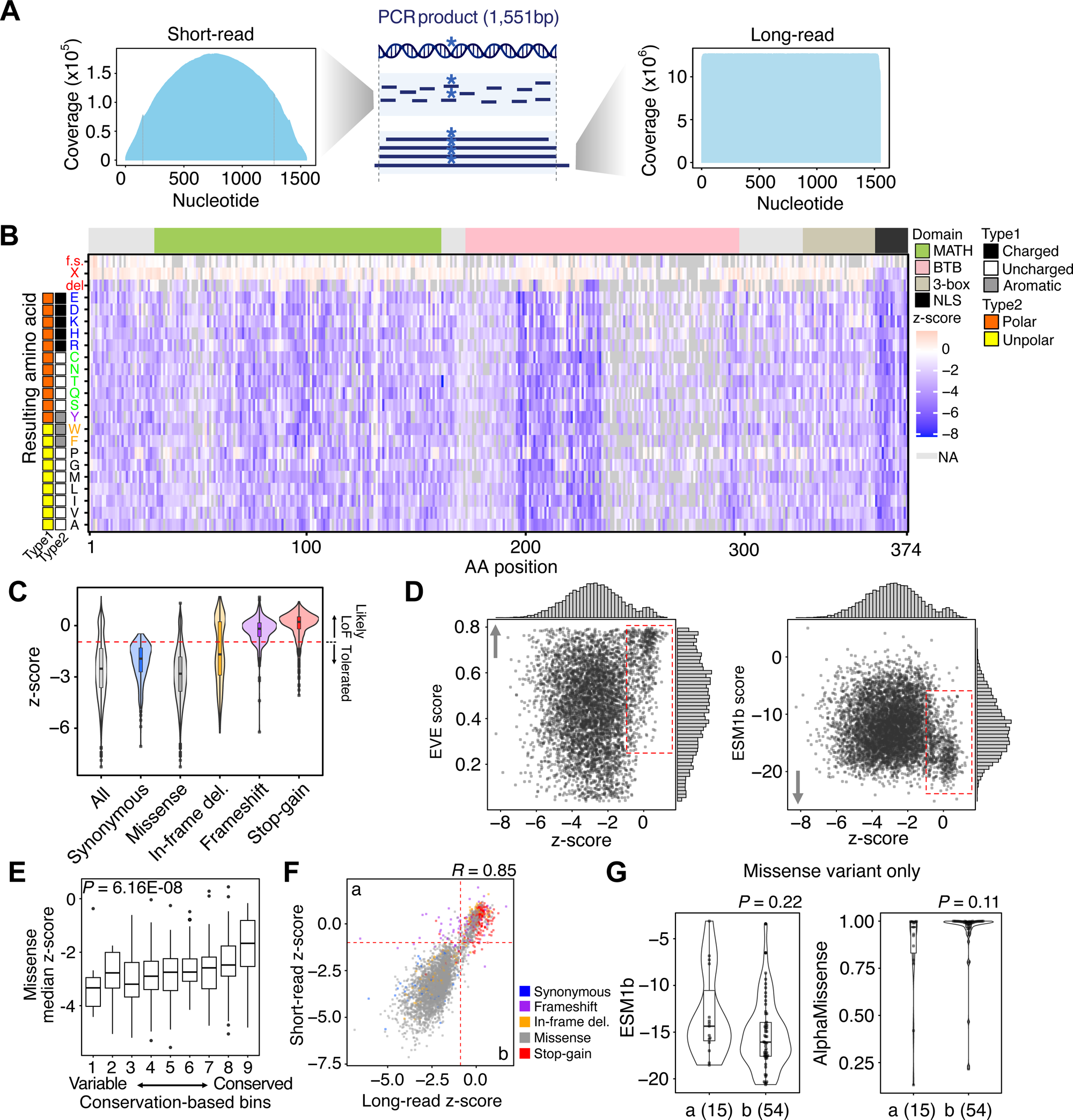
Generation of combined z-scores for the variant functional assessment. (A) Read coverage distributions of the reference template for short-read (left) and long-read (right) sequencing, and schematic illustration of read coverage patterns from each platform (center). Short-read sequencing showed a smooth, bell-shaped coverage profile across the ∼1.5 kb amplicon, with reduced depth toward the 5′ and 3′ ends consistent with fragmentation during library preparation. In contrast, long-read sequencing exhibited a relatively uniform coverage profile across the full amplicon, indicating even amplicon representation. (B) Heatmap showing combined z-scores for the covered *SPOP* variants. The x-axis represents the residue number and the y-axis the alternative amino acid. SPOP functional domains are indicated along the top. Amino acids are color-coded based on their biochemical properties. (C) Violin plot displaying z-score distribution by variant impact. The cutoff of ‘Likely LoF’ is represented as red dashed line. (D) Scatter plot of EVE and ESM1b scores for variants. ‘Likely LoF’ variants are depicted in the red dotted rectangles. Arrows beside panels indicate the direction of increasing pathogenicity. (E) Functionality score vs. evolutionary conservation. Median missense variant z-score values per each residue were used for comparison with conservation score in the residue level. *P*-value denotes the significance of the trend in a linear regression (F) Scatter plot showing correlation of z-scores from short-read and long- and short-read data. (G) ESM1b and AlphaMissense scores of variants classified as ‘Likely LoF’ by a single sequencing platform. ‘a’ and ‘b’ represent variants exclusively assigned as ‘Likely LoF’ from short- and long-read platforms, respectively. Two-tailed t-test *P*-values are shown.

**Figure 3.**
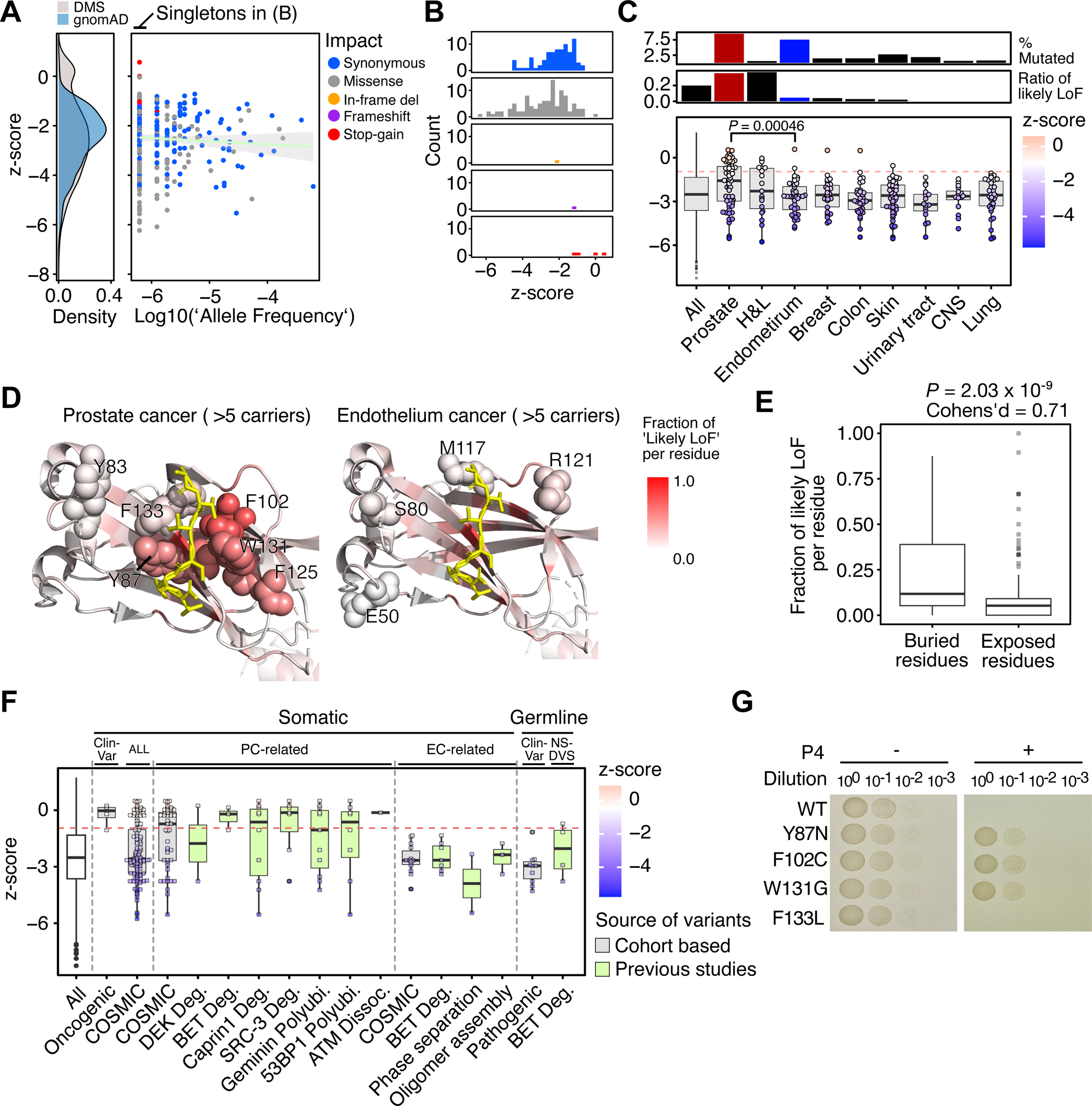
Validation of SPOP fitness scores. (A) Density plot of observed DMS variants and gnomAD variants (left). Scatter plot of z-scores with allele frequency in gnomAD (right). Variants on the far left part of the right panel are singletons. Variant impacts are distinguished by dot color; the linear regression line based on Log10 transformed allele frequency and z-score is indicated in light green. (B) Z-score distribution of gnomAD singleton variants according to impact on *SPOP*. (C) Z-score distribution of COSMIC variants according to primary tumor sites. The top panel indicates the percentage of patients with *SPOP* somatic mutations; the middle two panels the proportion of variants assigned as ‘Likely LoF’; and the bottom panel the distribution of z-scores per tumor site. H&L, Hematopoietic and lymphoid. CNS, Central nervous system. (D) Structural placement diagrams of recurrent variant residues (count > 5) found in prostate and endometrial cancers. Residues of interest were shown as color-coded spheres by fraction of ‘Likely LoF’ varaint per residue (PDB ID: 3HQI). Only the MATH domain and the SBC motf are displayed, represented as cartoons and yellow sticks, respectively. (E) Comparison of the fraction of ‘Likely LoF’ variants per residue value between buried or exposed regions. (F) Boxplots displaying the z-scores of variants reported in public databases or validated in previous studies. The plots are color-coded by source: grey for cohort-based data and green for variants from previous studies. (G) Yeast growth assay of variants associated with prostate cancer. Abbreviations: Deg., degradation; Polyubi., polyubiquitylation; Dissoc., dissociation.

**Figure 4.**
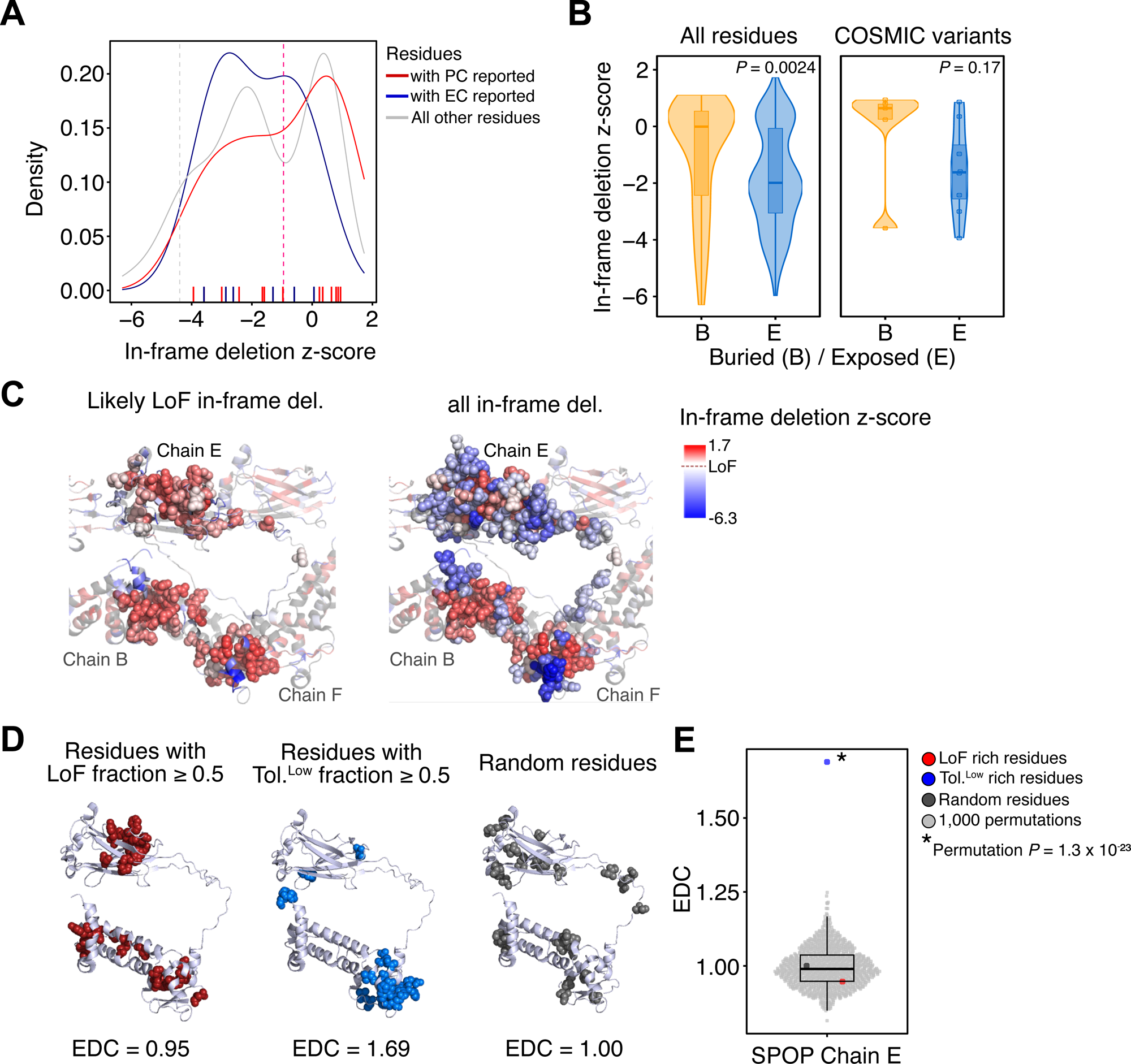
Interpretation of variant impact based on amino acid biochemical properties and structural context. (A) Dissection of COSMIC-overlapping residues for their z-score of the corresponding in-frame deletion on that position. (B) Stratification of in-frame deletion z-scores by residue solvent accessibility. (C) Residues with in-frame deletions are displayed as color-coded spheres on the oligomeric structure of SPOP. Colors indicate z-scores of in-frame deletions and “Chain” refers to SPOP monomer. (D) Residues exclusively enriched by variants assigned as ‘Likely LoF’ (left), ‘Tolerated’ variants with low scores (Tol.^Low^; middle), and randomly selected residues (right) were depicted in red, navy, and grey spheres, respectively. These sets of residues were used to calculate the EDC values. (E) EDC values from 1,000 permutations were shown as light dots with overlaid boxplot. EDC values corresponding to the residue groups in (D) were highlighted as larger colored dots (red, navy, and dark grey). Protein structure in (C-E) is based on the PBD ID: 8DWV and only Chain E was used for (D-E).

### Calculating extent of disease clustering (EDC)

EDC for the residues of interest was determined described in a previous study (61) with Chain E of the 8DWV PDB structure as reference. Residues were classified as disease residues if the fraction of variants at that position that were either ‘Likely LoF’ or ‘Tolerated’ with extreme lower-end z-scores was ≥ 0.5. Variants in the extreme low tail of the ‘Tolerated’ distribution were defined as those with a z-score below the 5th percentile of the synonymous variants distribution, using synonymous variants as an empirical reference. Randomly selected residues (*n* = 26, matching the size of disease residues) served as the negative control. Statistical significance was assessed over 1,000 permutations in which randomly selected sets were taken as disease residues and the remainder as non-disease residues.

## Results

### Measuring biological effect of SPOP in yeast

To dissect the function of *SPOP* at the amino-acid level, we delivered the DMS library into yeast, induced transgene expression by addition of progesterone, and determined *SPOP* sequences in surviving cells using short-read and long-read sequencing (Fig. 1A). Our library construction and delivery strategy includes insertion of a landing pad into the yeast genome to facilitate insertion of the library, which was made via concatenation of a mutation-containing tile (54-122 bp long) with the remaining 5’ and 3’ parts of the *SPOP* cDNA (SI Appendix, Fig. S1 and Methods).

We observed that expression of wild-type *SPOP* inhibits yeast growth (Fig. 1B), whereas depletion of specific functional regions within wild-type SPOP, namely the MATH, BTB, and 3- box domains (SI Appendix, Fig. S2), rescues it. To elucidate the molecular mechanism underlying this SPOP-induced growth inhibition, we performed tandem mass tag-labeled liquid chromatography-mass spectrometry and identified differentially expressed proteins in yeast cells with and without wild-type *SPOP* overexpression. A total of 100 DEPs were detected, 74 up- and 26 down-regulated (Fig. 1C, Dataset S3 and Fig. S4). To identify direct downstream targets of SPOP’s engagement in ubiquitination, we intersected the down-regulated proteins with SPOP-binding consequence (SBC) sequences. This revealed two proteins involved in the glycolysis pathway (FBA1 and CDC19) (Fig. 1C. D, SI Appendix, Fig. S4) reduced by the addition of *SPOP*. Notably, FBA1 encodes fructose-bisphosphate aldolase and catalyzes a reversible reaction for both glycolysis and glyconeogenesis (62); its impacts on cell growth and proliferation can be attributed to its role in glycolysis (63,64) and its regulation of the polymerase III RNA complex (65). Meanwhile, CDC19 encodes pyruvate kinase, which converts phosphoenolpyruvate to pyruvate and thereby finishes the glycolysis process in a partially compensatory manner; its depletion leads to growth arrest by impairing ATP generation (66–68). Notably, additional six out of 26 downregulated proteins are essential components of the glycolysis pathway (Fig.1D). Since the overexpression of CDC19 and FBA1 alone failed to rescue the lethal trait conferred by *SPOP* expression (SI Appendix, Fig. S6), it is possible that overall metabolic burden may have initiated the cell growth in our *SPOP*-expression system.

### Leveraging short- and long-read platforms for generation of a combined score

Many recent DMS studies have leveraged barcoding strategies combined with long-read sequencing to directly link mutations with their functional outcomes (69–71), while others have relied on short-read approaches that enable cost-efficient, high-depth quantification (11,15,72). However, library construction and sequencing can introduce additional unintended mutations that may not be fully captured by barcode-based designs alone. To take advantage of the complementary strengths of both platforms, we analyze our mutant libraries using short-read and long-read sequencing (Fig. 2A, SI Appendix, Table S1). The enrichment ratio of variants in the *SPOP*-expressed pool relative to the non-expressed pool was calculated separately for each platform (see Methods, Dataset S4). Two biological replicates were generated, and exhibited high correlation across treatment conditions and sequencing platforms (SI Appendix, Fig. S7).

To fully leverage the results from both sequencing platforms, we generated combined scores using inverse-variance weighted average values (Dataset S4). This led to the list of variants, including synonymous variants (designed = 374, observed = 354), missense-causing variants (designed = 7,106, observed = 6,940), stop-gain variants (designed = 374, observed = 369), single amino acid-deletion variants (designed = 374, observed = 270), and unintentionally occurred frameshift (*n* = 3,439) and in-frame (*n* = 3,099) variants (Fig. 2B, SI Appendix, Table S2).

Next, we stratified variants into two groups–‘Likely Loss-of-Function (LoF)’ and ‘Tolerated’– based on log2 fold changes and Benjamini-Hochberg adjusted *P*-values. Since the variant assessment assay is based on the yeast survival and functional *SPOP* variants are expected to reduce cell viability, we defined variants showing insignificant changes in viability as ‘Likely LoF’ (Log2FC > -1 and FDR > 0.1) and all remaining variants as ’Tolerated’. Then we assigned a single cutoff from the intersection of the z-score distributions of the ‘Likely LoF’ versus ‘Tolerated’ variant groups (Fig, 2C). This classification strategy demonstrated good concordance with predicted variant impact (Fig. 2C, SI Appendix, Fig. S8). For example, 88.7% of stop-gain and 85.7% of frameshift mutations were enriched in the ‘Likely LoF’ classification. Missense-causing variants and in-frame deletions exhibited a bimodal distribution with subsets enriched either across the ‘Likely LoF’ and ‘Tolerated’ classifications. Lastly, 90.7% of synonymous mutations were enriched in ‘Tolerated’ classification. To account for the possibility that these changes in SPOP function may be due to differences in expression rather than protein function, bulk RNA sequencing was performed to document *SPOP* mRNA expression. This sequencing revealed most variants to exhibit moderate changes in expression, and the correlation of expression with z-score was minimal (adjusted-*R^2^* = 0.009, SI Appendix, Fig. S9). Indeed, only 4.5% of the ‘Likely LoF’ variants (4 out of 89) statistically downregulated *SPOP* expression.

To better characterize variants classified as ‘Likely LoF’, we annotated each variant with EVE and ESM1b scores and observed that the ‘Likely LoF’ set was shifted toward more deleterious predictions compared to other variants (Fig. 2D). This finding suggests that variants in the ‘Likely LoF’ classification are likely to confer functional effects distinct from those of ‘Tolerated’ variants.

In addition, when we overlaid the evolutionary conservation scores of each amino acid residue from ConSurfDB with the median of missense-causing variant scores (73), we found significant association between median missense z-scores and conservation grades (*P*-value = 6.2 x 10^-8^ from linear regression, Fig. 2E).

When we compared scores derived from each sequencing platform, short- and long-read data showed strong concordance (Fig. 2F, Pearson’s correlation *R* = 0.85). Although a subset of variants were classified as ‘Likely LoF’ in a platform-specific manner (grid a and b in Fig. 2F; SI Appendix, Fig. S10), this discrepancy was not explained by differences in predicted deleteriousness: among missense variants, platform-specific calls showed no significant differences in matched ESM1b and AlphaMissense scores (Fig. 2G; Two-tailed t-test *P* = 0.22 and 0.11, respectively). Overall, while a subset of variants showed platform-specific ‘Likely LoF’ calls, we did not observe systematic biological differences. All told, our scoring model classified 713 out of 6,261 missense variants as ‘Likely LoF’.

### Validation of SPOP fitness scores

To further validate if our z-scores accurately represent the functionality of individual variants, we cross-referenced our scores with external databases and performed experiments to test the pathogenicity of selected variants. Since common variants tend to be benign, we first determined z-scores for the common missense variants listed in gnomAD; of these, 96.7% (58 out of 60) classified as ‘Tolerated’ (Fig. 3A-B). Similarly categorizing variants from tumor primary sites revealed that distributions of z-scores from the prostate and endometrium cancers differ, with variants from prostate cancer showing higher scores than those from endometrium cancer (two-sided *t*-test *P* = 4.6 x 10^-4^, Cohen’s *d* = 0.68) (Fig. 3C and SI Appendix, Fig. S11). This distinction was also observed when plotting the COSMIC variants from each tumor by frequency (SI Appendix, Fig. S12). To further compare prostate- and endometrial cancer-associated variants, we visualized recurrently mutated residues (count > 5) on the 3D structure of the MATH domain (Fig. 3D). Five of the six prostate-enriched residues localized near the SPOP- binding consensus (SBC) motif, and four of these five residues showed a high fraction (≥ 0.5) of ‘Likely LoF’ variants at that position. In contrast, recurrent endometrial cancer-associated residues showed lower ‘Likely LoF’ fractions (< 0.5) and were predominantly located at the MATH-domain interaction interface. These patterns are concordant with our ‘Likely LoF’ classification and with prior mechanistic models in which prostate cancer-associated *SPOP* mutations are predominantly LoF, whereas endometrial cancer-associated mutations exhibit more GoF-like properties, and they are consistent with the observed structural localization.

To define which classes of pathogenic variants are captured by our DMS readout, we examined which structural implications ‘Likely LoF’ variants have and where previously reported and experimentally validated variants fall in our score. We first assessed the structural distribution of ‘Likely LoF’ calls by collapsing variants into residues. Residues in structurally buried regions harbored a significantly higher fraction of ‘Likely LoF’ variants than solvent exposed residues (t- test *P* = 2.0 x 10^-9^ and Cohen’s *d* = 0.71; Fig. 3E and SI Appendix, Fig. S13), consistent with sensitivity to variants that compromise folding/stability. Next, we performed a comprehensive analysis integrating databases with clinical relevance (ClinVar and COSMIC) and experimentally validated variants (Fig. 3F). ClinVar-annotated somatic pathogenic variants and recurrent prostate cancer-associated variants from COSMIC were shifted toward the ‘Likely LoF’ range relative to endometrial cancer-associated variants (grey boxes in Fig. 3F). Consistent with this, variants known to interact with oncogenic proteins or proteins responsible for genomic stability (*e.g.*, BET (33), Caprin1 (74), SRC-3 (27), Geminin (75), and 53BP1 (30)) generally scored as ‘Likely LoF’, whereas validated endometrial cancer variants with GoF-like properties (enhanced polyubiquitination (33) and altered oligomer assembly (25)/phase separation (31)) were predominantly classified as ‘Tolerated’ (green boxes in Fig. 3F). In contrast, germline pathogenic variants showed weaker concordance with the ‘Likely LoF’ classification, including ClinVar variants and *de novo* variants associated with Nabais Sa-de Vries syndrome (NSDVS).

Furthermore, we evaluated the performance of our combined score model in distinguishing pathogenic variants from benign variants. Given the absence of a ClinGen Variant Curation Expert Panel (VCEP) for *SPOP*, we used ClinVar as the primary source of previously reported variants. ClinVar variants annotated as *Pathogenic/Likely Pathogenic* were used to define a reference pathogenic set and were stratified into ClinVar-Somatic and ClinVar-Germline, with gnomAD missense variants used as a benign reference (SI Appendix, Table S3). Our scoring system achieved the highest selection for the ‘ClinVar-Somatic’ set (LR+ = 22.5). Given the limited size of the ‘ClinVar-Somatic’ set (*n* = 4), we expanded somatic pathogenic set with recurrent (count >1) prostate cancer-associated COSMIC variants (COSMIC-PC), which retained robust performance (LR+ = 16.94). In contrast, ‘ClinVar-Germline’ showed poor concordance with our LoF-based assay readout (LR+ = 0), whereas the curated germline subset associated with Nabais Sa-de Vries syndrome and proposed LoF mechanisms showed slightly improved performance (LR+ = 7.5). This suggests that our scoring system, which captures the SPOP-mediated degradation of essential proteins of yeast growth, specifically identifies variants leading to LoF via interaction defects and structural destabilization. While this robustly predicts the pathogenicity of somatic variants driving prostate cancer, it shows limited sensitivity towards germline mutations.

Next, we experimentally tested variants found in high frequencies among the prostate cancer patients through assessing yeast survival (Fig. 3G). As expected, variants classified as ‘Likely LoF’ (*i.e*., p.Y87N, p.F102C, and p.W131G) failed to arrest yeast growth, whereas ‘Tolerated’ variants that were annotated as benign, found in endometrial cancers, and other variants from prostate cancers (*e.g.*, Y123C and F133L) maintained the growth inhibition phenotype (Fig. 3G and SI Appendix, Fig. S14).

### Utility of in-frame deletion scores

The addition of in-frame deletion variants allowed us to distinguish whether a given amino acid residue functions as LoF or GoF, because we posited that GoF residues would not result in any functional change if removed, whereas LoF residues might. Plotting the z-score values for amino acid residues with in-frame deletion revealed that the half of the residues relevant to prostate cancer (6/13) were enriched in the ‘Likely LoF’ classification for in-frame deletion (Fisher’s exact test *P* = 3.3 x 10^-3^) (Fig. 4A). Meanwhile, most residues recurrently observed in endometrium cancer (4/6) were categorized into the ‘Tolerated’ classification for in-frame deletion-based z-scores.

To elucidate tolerance to in-frame deletion, we aligned solvent accessibility information with residues and observed that the in-frame deletion-based z-score differed significantly depending on whether a residue was buried or exposed (two-sided t-test *P* = 2.4 x 10^-3^, Cohen’s *d* = 0.35). Specifically, residues located in exposed sites were more likely to tolerate in-frame deletions (Fig. 4B). Previous studies have reported that residues impacted by variants recurrently observed in prostate and endometrium cancers exhibit distinct structural distributions: those related to prostate cancer are predominantly enriched in the substrate-binding site of the MATH domain, whereas those related to endometrium cancer are more enriched at SPOP domain interfaces that are important for SPOP self-assembly (25,33). This structural distinction has been exploited to explain the divergent pathways by which *SPOP* mutations promote tumorigenesis in those two cancer types (25,31). In our 3D protein structure annotated with the z-score of in-frame deletions as color of spheres (Fig. 4C), residues with ‘Likely LoF’ z-scores are buried inside by those classified as ‘Tolerated’. Ultimately, inference from this analysis led us to pinpoint a group of variants likely to confer GoF inferred from the in-frame deletion z-scores being found most frequently in the BTB domain, followed by the MATH domain (SI Appendix, Fig. S15).

We also investigated the intrinsic importance of the reference amino acids, in which the most pronounced change in z-scores was observed upon alteration of tryptophan (SI Appendix, Fig. S16A). While most reference amino acids exhibited ‘Intermediate’ mean z-scores upon substitution, replacing tryptophan generally resulted in mean z-scores associated with ‘Possible LoF’, except for substitutions on a few amino acids with nonpolar side chains (SI Appendix, Fig. S16B).

Disease-causing variants that operate by different molecular mechanisms are known to exhibit distinct tendencies with regard to clustering on 3D protein structure (61). To assess whether our DMS model reflects this phenomenon, we extracted residues that are enriched with ‘Likely LoF’ variants and variants at the other end of the z-score distribution. This assignment is based on our assumption that GoF variants would be enriched in the low end of the z-score distribution (SI Appendix, Fig. S17). Extent of disease clustering (EDC) values of residues enriched with ‘Tolerated’ variants with low z-scores showed an EDC of 1.69, whereas both those associated with ‘Likely LoF’ variants and randomly selected residues exhibited EDC values close to 1 (Fig. 4D; see Methods). Upon 1,000 permutations of randomly selected residues, the EDC observed for residues associated with low z-scores was significantly higher (permutation-based *P* = 1.3 x 10^-23^, Fig. 4E). This observation indicates that ‘Tolerated’ variants with extreme z-scores might have additional functionality compared to other ‘Tolerated’ variants.

## Discussion

*SPOP* possesses a remarkable context-dependent functionality in different tumor types. In particular, mutations recurrently observed in prostate cancer cause the gene to act as a tumor-suppressor, but mutations observed in endometrial cancer appear to confer a GoF-like effect. Also, somatic mutations occur widely throughout the gene, making its functional annotation and clinical implication challenging. Despite active studies on the tumorigenesis of SPOP, only variants located in well-characterized structural regions have been reported, and a database categorizing pathogenic variants is lacking.

To evaluate the function of each variant at a higher resolution, we performed a DMS targeting SPOP, covering 96.4% of its all possible variants. Using yeast cell survival as a functional indicator, we were able to resolve variants into binary classification: ‘Likely LoF’ and ‘Tolerated’. Most stop-gain and frameshift variants fell into the ‘Likely LoF’, and synonymous variants into the ‘Tolerated’; missense variants and in-frame deletions showed wider classification distributions. Notably, our DMS scores classified the majority of gnomAD variants into the ‘Tolerated’ classification and differential score distributions were evident between variants recurrently observed in prostate and in endometrium cancer.

In addition to the functional dissection of *SPOP*, we demonstrate two technical improvements over previous DMS studies. First, we attempted to utilize long reads to fully cover the mutation profile of the DMS library. Albeit the long-read sequencing allowed us to document additional inadvertent mutations in the non-targeted portion of the gene, the fitness scores based on short-read-only and combined sets did not differ substantially in overall concordance rate. Second, we created in-frame deletions across most amino acid residues, which provided indications of whether a given residue would function in a GoF or LoF manner. In particular, integration of the positional enrichments of these variants provided molecular evidence toward the determination of GoF variants, predominantly found in endometrium cancer patients.

Our comparisons against known variants define the scope of our scoring system for SPOP. We observed that curated somatic driver variant sets were enriched in the ‘Likely LoF’ class, and literature-curated variants with experimentally validated LoF mechanisms similarly clustered within the ‘Likely LoF’ range. In contrast, endometrial cancer-associated variants were preferentially assigned outside the ‘Likely LoF’ range, consistent with the previous reports that many of these mutations exhibit GoF-like properties. Accordingly, our score showed robust performance when evaluated using the ‘ClinVar-Somatic’ and ‘COSMIC-PC’ reference set (LR+ = 22.5 and 16.94, respectively). Together, these results support that our assay reliably assigns variants as ‘Likely LoF’ when they impair SPOP function in a manner consistent with loss of substrate recognition and/or degradation activity.

Multiple DMS studies have been performed in the cancer context to categorize tumorigenesis variants and establish scoring models based on cellular fitness readouts (76–79). The assay strategy often includes cell abundance (12), homology-directed repair reporter gene activity (80), and physical interaction with a partner protein (81). Our DMS library was synthesized through tile-based cloning, whereas several studies introduced mutations via CRISPR-based editing (82,83). While tile-based cloning allows saturation for all kinds of mutations, the CRISPR-based method allows comparison of the scoring system with nucleotide-level cancer signature analysis (77) and impacts on splicing or untranslated regions (78).

Our study has several limitations. The assay system is yeast-based and may not cover all substrate interactions that occur in human cells. Yeast lacks an endogenous *SPOP* ortholog and does not encode recognizable homologs of major mammalian SPOP substrates (such as ERG (84,85), TRIM24 (86), c-MYC (87), NANOG (88), PD-L1 (89), IRF1 (90), SRC-3 (27), DAXX (91), DEK (92), GLI2 (93), and AR (94)). However, yeast encodes a diverse repertoire of SCF/F-box proteins, which operate as substrate adaptors in an analogous manner to mammalian substrate-recognition modules. Moreover, the core elements of the ubiquitin-proteasome system required for SPOP-mediated ubiquitination are evolutionarily conserved. Yeast expresses orthologues of Cullin3 (Cdc53), Rbx1 (Hrt1), and multiple E2 ubiquitin-conjugating enzymes (including Ubc4/Ubc5), which together form a functional CUL3-type Cullin-RING ubiquitin ligase (CRL) scaffold (SI Appendix, Fig. S3). Collectively, these conserved components provide a mechanistically compatible cellular environment that allows heterologous human SPOP to engage the ubiquitination machinery despite the absence of its native substrates.

Secondly, it would be also possible to use the library to assay against a specific substrate, oligomerization status, or formation of phase separation of SPOP (31,95). However, our assay also showed limitations in distinguishing neutral variants from GoF variants, as endometrial-associated variants enriched in ‘Tolerated’ classification. This likely reflects that the ‘Tolerated’ class contains a mixture of truly neutral variants and pathogenic variants acting through non-LoF mechanisms, which in turn reduces the interpretability of negative calls and contributes to weaker negative likelihood ratios (SI Appendix, Table 3). Therefore, ‘Tolerated’ calls should be interpreted with caution and may not be used as a filter for excluding potentially relevant variants.

Lastly, when we evaluated the performance of our scoring system in predicting pathogenicity, we found that it does not fully capture the malfunctionality of SPOP caused by germline mutations. This discrepancy may reflect distinct underlying molecular pathway in somatic versus germline contexts, which are associated with different disease symptoms (*i.e.*, somatic mutations cause tumors, while germline mutations cause neurodevelopmental disorder; OMIM #618828 and #618829).

As told, this work presents a nearly complete functional atlas of SPOP single amino acid variants that enables refined interpretation of their potential clinical impact. This resource may provide a foundation for future mechanistic and translational studies through linking variant- specific functional consequences to tumor biology and therapeutic strategies.

## Acknowledgment

We thank Theragen Etex and SNUH proteomics facility for technical supports and Eun Ha Kim for administrative support.

## Author contributions

Seong Kyoon Park: Data curation, Investigation, Methodology, Resources, Validation, Writing – original draft Jeongha Lee: Data curation, Investigation, Formal analysis, Methodology, Software, Validation, Visualization, Writing – original draft Seon Ju Park: Methodology, Writing – review & editing Ye Na Kim: Methodology, Project administration, Writing – review & editing Gi Hyun Shin: Methodology, Writing – review & editing Kisoon Dan: Formal analysis, Writing – review & editing Hee-Jung Choi: Methodology, Software, Supervision, Writing – review & editing Dohyun Han: Methodology, Resources, Supervision, Writing – review & editing Byung Joon Hwang: Conceptualization, Funding acquisition, Methodology, Supervision, Writing – original draft Murim Choi: Conceptualization, Funding acquisition, Methodology, Supervision, Writing – original draft

## Data availability

All sequencing data were deposited at Short Read Archives BioProject PRJNA1304161, individual variant scores are available at MaveDB (ID: urn:mavedb:00001258-a-1), and Mass spectrum based based proteomics data were deposited at PRIDE (ID: PXD067557, will be accessible upon publication). All codes for calculating scores in DMS models are available on Zenodo at https://doi.org/10.5281/zenodo.16787854.

