## Supplemental materials for "Functional dissection of *SPOP* on the amino acid level reveals a comprehensive functional landscape of variants during tumorigenesis"

**Contents:**

**Supplementary Table 1.** Summary table of short- and long- read sequencing reads.

**Supplementary Table 2.** Number of expected/observed variants by their impact.

**Supplementary Table 3.** Predictive likelihood ratio for *SPOP* variants called as pathogenic and benign.

**Supplementary Figure 1.** Vector maps of the landing pad and DMS library.

**Supplementary Figure 2.** Yeast growth assay of *SPOP* with deletions of various domains.

**Supplementary Figure 3**. Expression of CUL3-type CRL and house-keeping genes (left) and proteins (right) in the yeast.

**Supplementary Figure 4**. Yeast GO terms from the profile of differential expressed proteins.

**Supplementary Figure 5.** Protein sequence of putative target protein of SPOP in yeast.

**Supplementary Figure 6**. Assessment of growth rescue by overexpression of putative SPOP substrates.

**Supplementary Figure 7**. Correlation between two biological replicates.

**Supplementary Figure 8**. Z-scores of variants by amino acid positions.

**Supplementary Figure 9.** Differential enrichment of reads from bulk-RNA sequencing between *SPOP*-expressed and non-expressed pool.

**Supplementary Figure 10.** Stratified scatter plot of figure 2F.

**Supplementary Figure 11.** Enrichment of variants reported in COSMIC by DMS z-score and the number of cases.

**Supplementary Figure 12**. Distribution of z-scores for recurrent variants reported in COSMIC.

**Supplementary Figure 13.** Positional enrichment of likely LoF variants in the SPOP structure.

**Supplementary Figure 14.** Validation of SPOP variants using small scale spotting assay.

**Supplementary Figure 15.** Proportion of in-frame deletion’s classification by SPOP domain

**Supplementary Figure 16.** Intrinsic characteristics of reference and altered amino acids related to z-score.

**Supplementary Figure 17.** Assessment of extreme subset of ‘Tolerated’ variants.

| **Sequencing platform** | **Short-reads** | | | | **Long-reads** | | | | | | | |
| --- | --- | --- | --- | --- | --- | --- | --- | --- | --- | --- | --- | --- |
| Sample | Non-expressed pool replicate 1 | SPOP-expressed pool replicate 1 | Non-expressed pool replicate 2 | SPOP-expressed pool replicate 2 | Non-expressed pool replicate 1 | | | SPOP-expressed pool replicate 1 | | | Non-expressed pool replicate 2 | SOP-expressed pool replicate 2 |
| Number of PASS reads* | 5.45E+07 | 5.34E+07 | 6.06E+07 | 6.97E+07 | 8.39E+06 | 1.13E+07 | 9.14E+06 | 1.42E+07 | 1.53E+07 | 9.18E+06 | 8.78E+06 | 8.20E+06 |
| Median length of reads | 300 | 300 | 300 | 300 | 2805 | 1590 | 1590 | 1590 | 1590 | 1590 | 1600 | 1590 |

**Supplement Table 1. Summary table of Nanopore sequencing reads.** *The number of reads was counted at tha paired-end level for short read based results.

| **Designed** | **Type** | **n_Expected** | **n_Observed_short** | **n_Observed_long** | **n_Observed_total** |
| --- | --- | --- | --- | --- | --- |
| Yes | Missense | 7106 | 6,936 | 5,834 | 6,940 |
| Yes | Stop-gain | 374 | 369 | 348 | 369 |
| Yes | In-frame deletion | 374 | 266 | 233 | 270 |
| Yes | Synonymous | 374 | 354 | 191 | 354 |
| No | Frameshift | 0 | 2,705 | 1,730 | 3,439 |
| No | In-frame insdel | 0 | 1,913 | 1,131 | 2,750 |
| No | In-frame ins | 0 | 325 | 43 | 349 |

**Supplement Table 2. Number of expected/observed variants by their impact.** Counts were extracted from raw variant count matrix. Column with prefix ‘_il’, ‘_np’, and ‘_total’ display counts from Illumina sequencing, Nanopore sequencing, and merged count, respectively.

| **Variant type** | **Variant set*** | **Number of variants (P/B)** | **Pathogenic variants** | | | **Benign variants** | | | **AUC** |
| --- | --- | --- | --- | --- | --- | --- | --- | --- | --- |
|  |  |  | **Tolerated** | **Likely LoF** | **LR** | **Tolerated** | **Likely LoF** | **LR** |  |
| Missense | ClinVar-Somatic | 4/60 | 1 | 3 | 22.5 | 58 | 2 | 0.26 | 0.94 |
|  | COSMIC-PC | 32/60 | 15 | 17 | 16.94 |  |  | 0.48 | 0.74 |
|  | ClinVar-Germline | 10/60 | 10 | 0 | 0 |  |  | 1.03 | 0.44 |
|  | Nabais Sa et al., LoF | 4/60 | 3 | 1 | 7.5 |  |  | 0.78 | 0.63 |
| Stop-gain | Stop-gain | 344/60 | 39 | 305 | 26.6 |  |  | 0.12 | 0.91 |

**Supplementary Table 3. Predictive likelihood ratio for *SPOP* variants called as pathogenic and benign.** *P, pathogenic. B, benign. LR, likelihood ratio. AUC, area under curve.

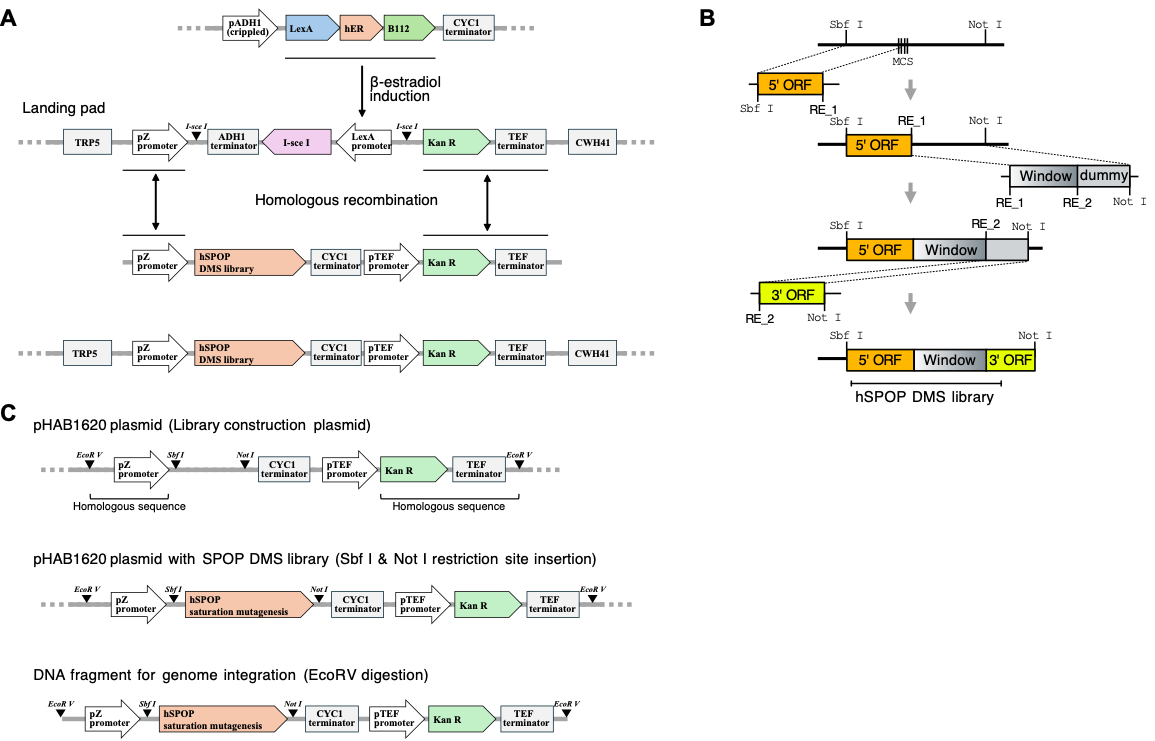

**Supplement figure 1. Vector maps of the landing pad and DMS library.** (A) The integration of the DMS library into the landing pad in the yeast genome is illustrated. The top construct displays the artificial transcription factor, which is activated by ß-estradiol induction and drives the activation of the LexA promoter in the landing pad. The constructs in the second and third rows represent the landing pad and DMS library, respectively. The sites of homologous recombination are indicated by double-headed arrows. The bottom construct represents the final result of the DMS library integrated into the landing pad. (B) Overview of the sequential steps in the DMS library cloning pipeline. ‘Tile’ represents synthesized oligonucleotides containing mutations, while 5’ ORF and 3’ ORF cover the remaining flanking sequences of ORF. (C) Cloning of the DMS library into a vector backbone compatible with the landing pad construct. The first row illustrates the vector backbone, the second row the vector after DMS library insertion, and the third the restricted form ready for yeast genome integration.

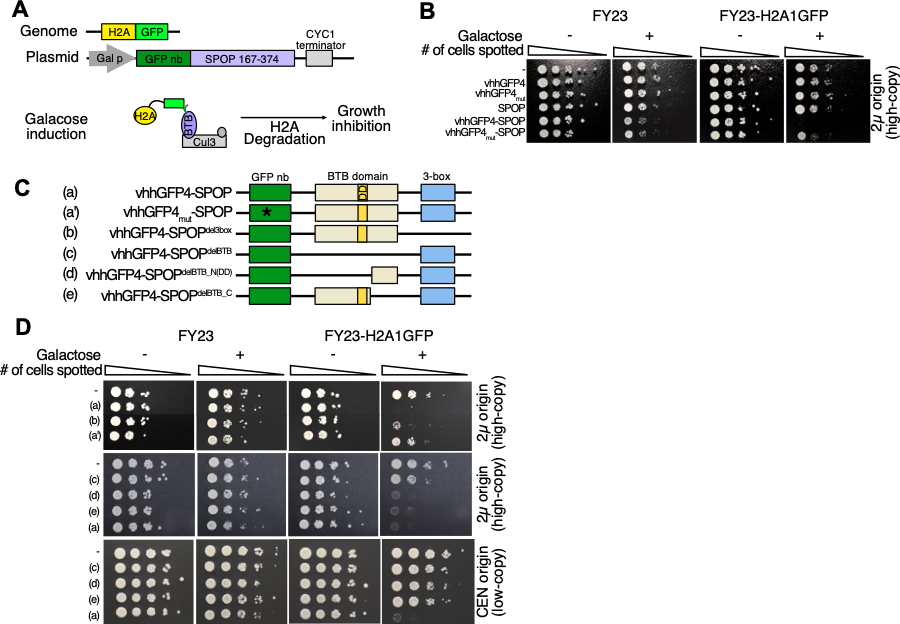

**Supplement figure 2. Yeast growth assay of SPOP with deletions of various domains**. (A) A *SPOP* construct in which the MATH domain was replaced with a GFP nanobody served as the template for generating mutations. A yeast strain was engineered to constitutively express H2A1 conjugated with GFP, which serves as a substrate for GFP nanobody and induces growth inhibition through degradation. ‘GFP nb’ refers to a GFP nanobody (vhhGFP4) that captures H2A-GFP by mimicking the role of the MATH domain. (B) displays yeast growth outcomes using constructs introduced in (A). (C) displays mutant clones in which each domain was individually removed, indicated by superscript labels. vhhGFP4 corresponds to the construct encoding the GFP nanobody alone, while vhhGFP4mut indicates the same construct with the vhhGFP4 with the CDR3 deleted. SPOP refers specifically to the 167-374 region. (D) shows individual yeast growth assays for known pathogenic mutations.

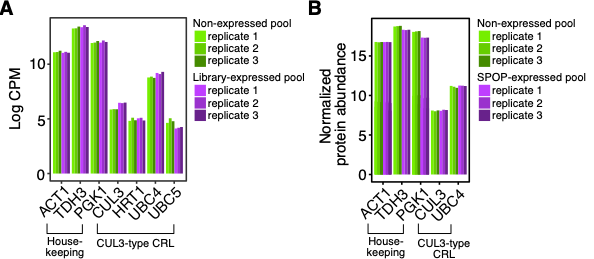

**Supplementary figure 3. Expression of CUL3-type CRL and house-keeping genes (left) and proteins (right) in the yeast.** (A) mRNA expression and (B) protein abundance of house-keeping genes/proteins and genes/proteins associated with the CUL3 type CRL scaffold. The ‘non-expressed pool’ denotes baseline samples lacking induction of the variant library or WT SPOP, whereas ‘library/SPOP-expressed pool’ denotes treated samples expression SPOP. Biological replicates are indicated by distinct colors.

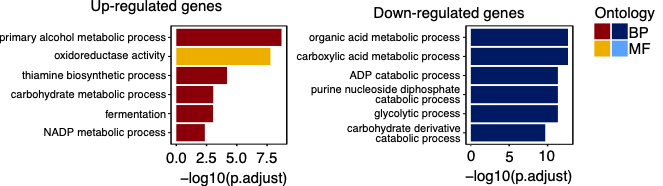

**Supplement figure 4. Yeast GO terms from the profile of differential expressed proteins.** Gene ontology terms of differentially expressed proteins were displayed with a bar plot. Terms associated with biological processes were highlighted with a darker color than those associated with molecular function.

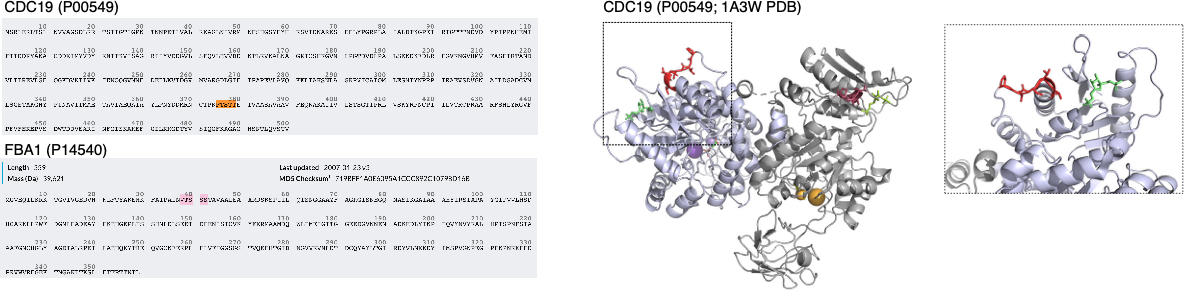

**Supplement figure 5. Protein sequence of putative target protein of SPOP in yeast.** The left panel with protein sequences of CDC19 and FBA1, extracted from UniProt, and their predicted SBC sequence were highlighted with orange color. The right panel displays the CDC19 structure (PDB ID: 1A3W). The SBC sequences were depicted with red color sticks.

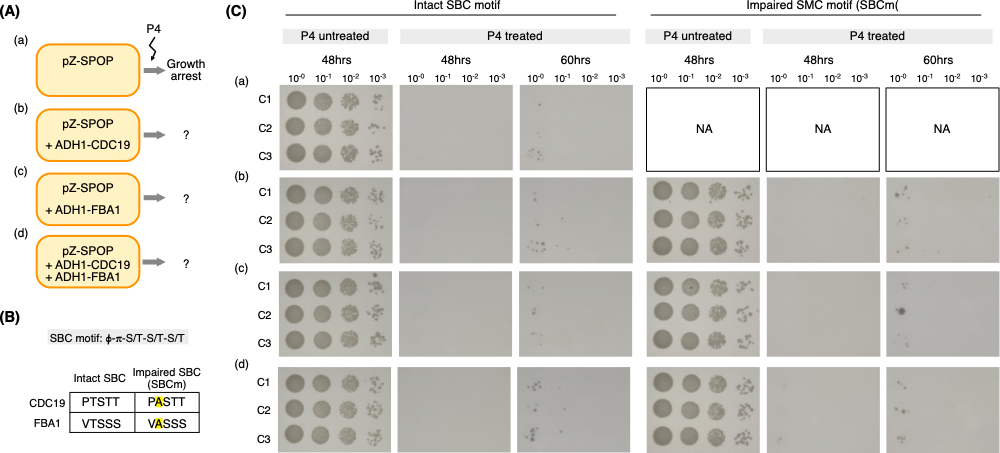

**Supplementary figure 6.** **Assessment of growth rescue by overexpression of putative SPOP substrates.** (A) Schematic representation of the rescue assay strategy. Yeast cells harboring the *SPOP* expression vector were co-transformed with vectors expressing CDC19, FBA1, and both. The genes are under the influence of constitutively active ADH1 promoter. (B) Introduction of SPOP-binding consensus (SBC) motif mutations to prevent potential degradation by SPOP (designated as CDC19-SBCm and FBA1-SBCm). (C) Serial spotting assay using the constructs described in (A) and (B). Yeast cultures were spotted onto plates with or without progesterone (P4) to induce *SPOP* expression.

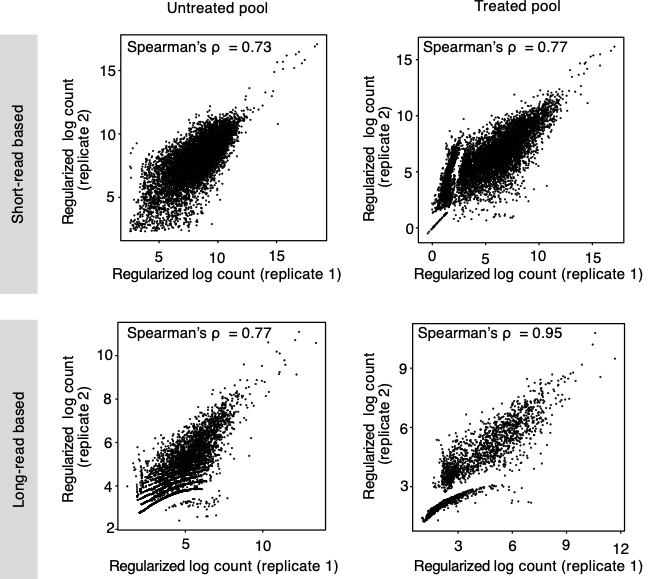

**Supplementary Figure 7**. **Correlation between two biological replicates.** Regularized log-transformed variant counts were used to confirm the correlation between the two biological replicates. The rlog function from DESeq2 was used to calculate the regularized log transform from DESeqDataSet. The top and bottom panels display data generated from short-read and long-read platforms, respectively. The left and right panels display data from untreated and treated pools, respectively. Replicate correlations (Spearman’s ρ) were 0.73 for the untreated pool (short-read), 0.77 for the treated pool (short-read), 0.77 for the untreated pool (long-read), and 0.95 for the treated pool (long-read).

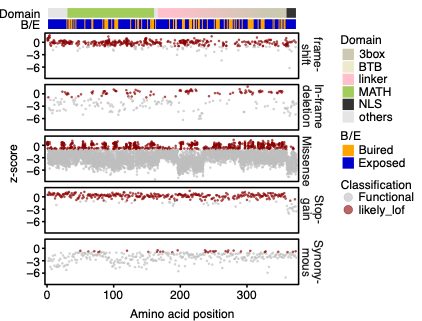

**Supplementary Figure 8**. **Z-scores of variants by amino acid positions.** Scatter plot displaying the z-scores of variants across the mutation impacts and positional enrichment of variants, highlighting tier-specific enrichment at certain positions.

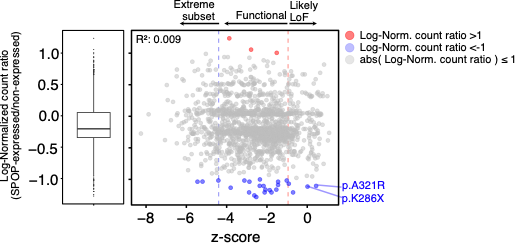

**Supplementary Figure 9. Differential enrichment of reads from bulk-RNA sequencing between *SPOP*-expressed and non-expressed pool.** The log-normalized count ratio between the SPOP-expressed versus non-expressed pools was shown as a boxplot (left). Variants detected in bulk RNA-sequencing were depicted according to their z-scores and log-normalized count ratios. The adjusted R squared between z-scores and log-normalized count ratios was 0.009. Variants with upregulated and downregulated expression were highlighted in red and navy, respectively.

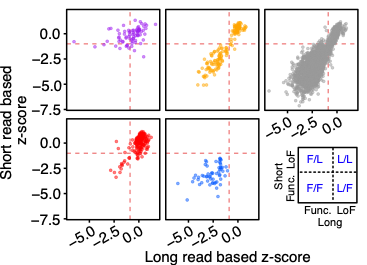

**Supplementary figure 10. Stratified scatter plot of Figure 2F.** Scores generated from the long-read platform showed improved concordance with the functional classification. Among frameshift variants, 23 were classified as ‘Likely LoF’ by short-read-derived scores but ‘Tolerated’ by long-read scores, whereas 3 were classified as ‘Tolerated’ by short-reads and ‘Likely LoF’ by long-read platform. For stop-gained variants, 4 variants were classified as ‘Likely LoF’ by the short-read platform and ‘Tolerated’ by long-reads, while 10 showed the opposite classification pattern.

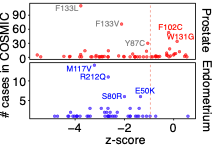

**Supplementary Figure 11. Enrichment of variants reported in COSMIC by DMS z-score and the number of cases.** Variants reported in prostate and endometrial cancers were displayed according to their z-scores and the number of reported cases. Red dashed line indicated the z-score thresholds used for likely LoF classification. Variants reported in prostate cancer with high z-scores were labeled with red, while those with low z-scores were labeled in grey. Variants reported in endometrial cancer were labeled in navy.

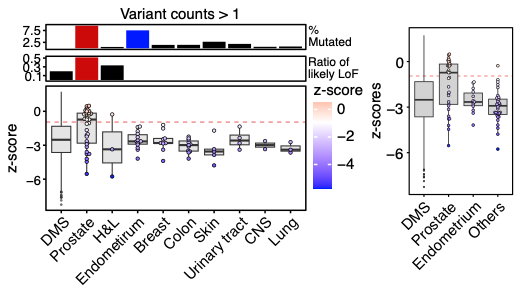

**Supplementary figure 12. Distribution of z-scores for recurrent variants reported in COSMIC (Figure 3C shows all COSMIC variants).** (Left) Distribution of z-scores for variants with a count >1. This panel corresponds to Figure 1C but filters out singleton variants to minimize sporadic passenger mutations. (Right) Aggregated variant scores by cancer types where all cancer types excluding prostate and endometrium are combined into a single ‘Others’ group.

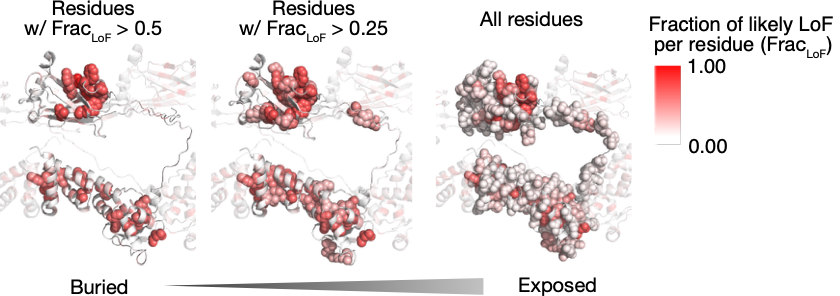

**Supplementary figure 13. Positional enrichment of likely LoF variants in the SPOP structure**. (A) Visualization of the SPOP structure derived from PDB 8DWV (Chain E) highlighting residues with high likely LoF fraction (Frac_LoF_ = In a given residue, number of variants that were called as likely LoF divided by number of all variants). The panels display residues at different inclusion thresholds: Frac_LoF_ > 0.5 (left), Frac_LoF_ > 0.25 (middle), and all residues (right). Residues are represented as colored spheres based on their Frac_LoF_ values.

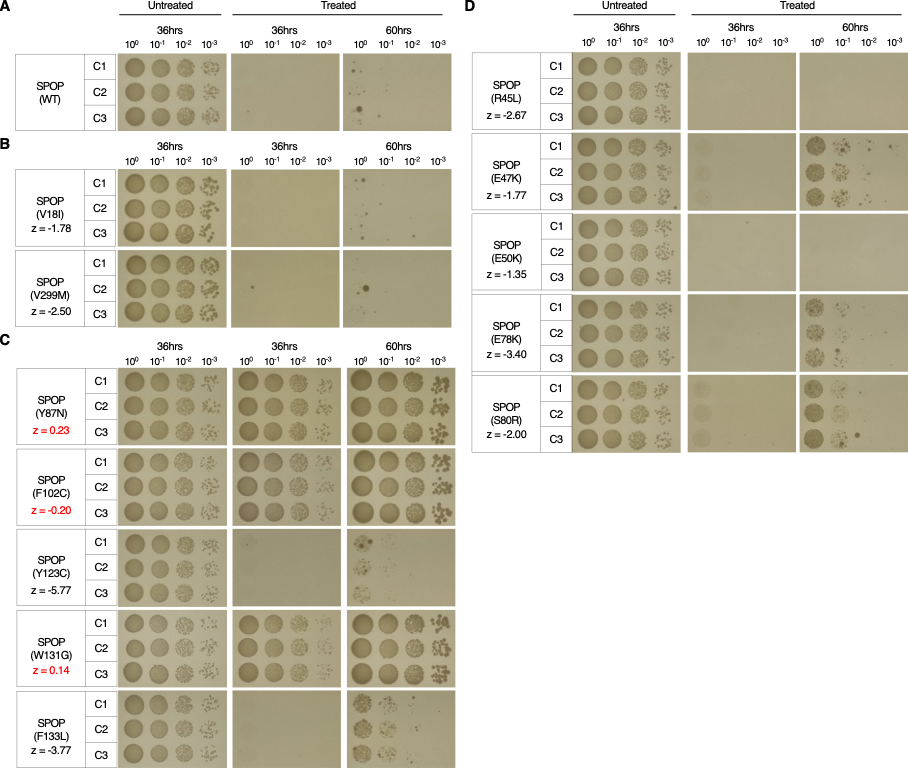

**Supplementary figure 14. Validation of *SPOP* variants using small scale spotting assay.** WT SPOP and indicated variants were stably integrated into the yeast genome. Cells were serially diluted and spotted onto plates containing 500 nM progesterone to induce protein expression. (A) WT SPOP exhibits strong cytotoxicity upon induction. (B) Benign variants, (C) prostate cancer, and (D) endometiral cancer-related variants exhibit cytotoxicity comparable to WT. Z-scores in red font denote values that passed the cutoff and are called likely LoF, which all showed loss-of-lethal trait.

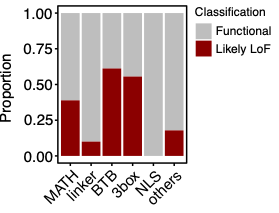

**Supplementary figure 15. Proportion of in-frame deletions’ classifications by SPOP domain.** Residues were stratified based on both their protein domain and the classifications assigned to their in-frame deletions.

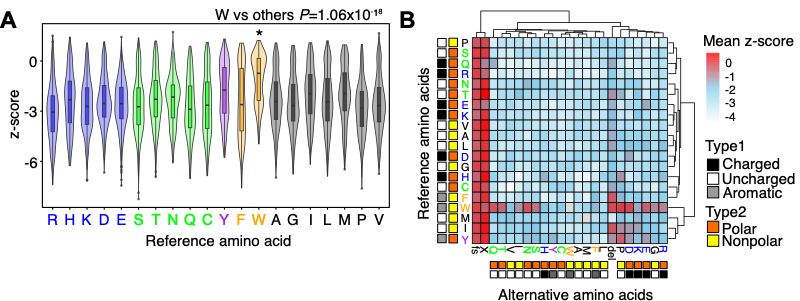

**Supplementary figure 16. Intrinsic characteristic of reference and altered amino acids related to z-score.** (A) Violin plot showing z-scores of variants grouped by reference amino acids, with colors indicating the intrinsic characteristics of each amino acid. Variants introduced on tryptophan (W) residues showed significantly different scores (*P* = 1.1 x 10^-18^). Color-codes of amino acids are as following: charged polar (blue), uncharged polar (green), aromatic polar (purple), aromatic nonpolar (orange), and uncharaged nonpolar (black). (B) Mean z-scores of reference amino acid and alternative amino acid pairs. Hierarchical clustering nodes were generated through an R package *pheatmap*.

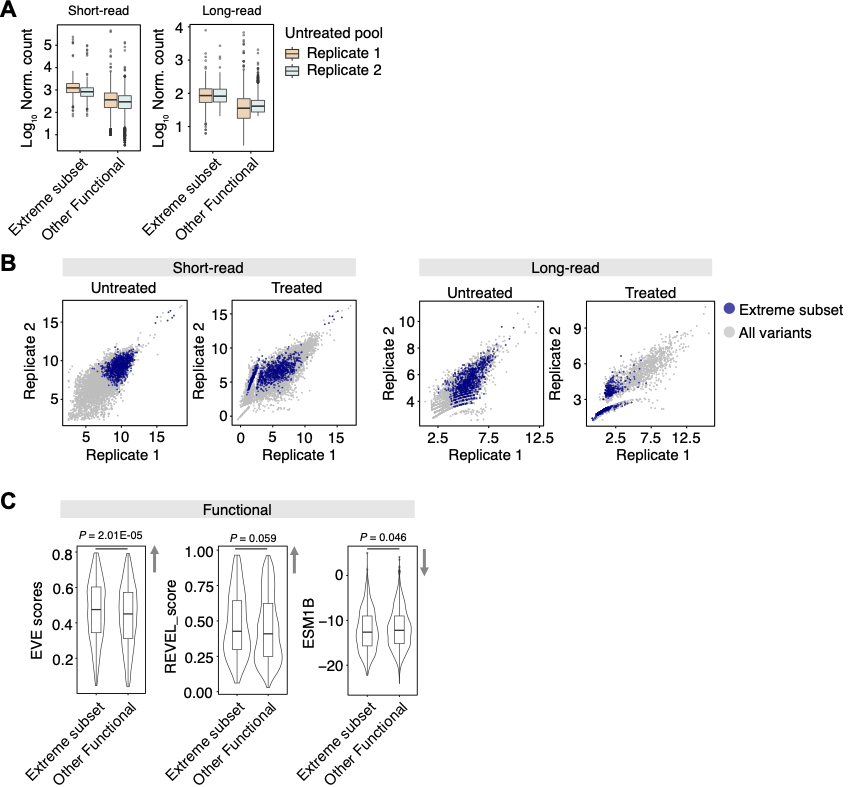

**Supplementary figure 17. Assessment of extreme subset of ‘Tolerated’** **variants.** (A) Distribution of log 10 transformed normalized counts for variants in the untreated pool, visualized as a boxplot. Variants were stratified into two groups: extreme subset, and other ‘Tolerated’. (B) Replicate consistency analysis using regularized log-transformed counts. Extreme subset was highlighted in blue to visualize their correlation between replicates. (C) Computational pathogenicity scores (EVE, REVEL, and ESM1b) across the three variant groups, displayed as violin plots. Arrows indicate the direction of increased deleteriousness. *P*-values were calculated using a two-tailed t-test comparing the extreme subset (*n* = 967) versus other ‘Tolerated’ (*n* = 5,137).
